# Neurodevelopmental deviations in schizophrenia: Evidences from multimodal connectome-based brain ages

**DOI:** 10.1101/2024.12.22.629311

**Authors:** Yun-Shuang Fan, Pengfie Yang, Yucheng Zhu, Wenfeng Jing, Yong Xu, Jing Guo, Wei Huang, Huafu Chen

**Affiliations:** The Clinical Hospital of Chengdu Brain Science Institute, School of Life Science and Technology, University of Electronic Science and Technology of China, Chengdu, China; Department of Psychiatry, First Hospital/First Clinical Medical College of Shanxi Medical University, Taiyuan, China; MOE Key Lab for Neuroinformation, High-Field Magnetic Resonance Brain Imaging Key Laboratory of Sichuan Province, University of Electronic Science and Technology of China, Chengdu, China

**Author notes:** Corresponding authors: Huafu Chen.

**Keywords:** *brain age*, *connectome*, *multimodal MRI*, *neurodevelopment*, *schizophrenia*

## Abstract

**Background:** Pathologic schizophrenia process originate early in brain development, leading to detectable brain alterations via structural and functional magnetic resonance imaging (MRI). Recent MRI studies have sought to characterize disease effects from a brain age perspective, but developmental deviations from the typical brain age trajectory in youths with schizophrenia remain unestablished.

**Aim:** This study investigated brain development deviations in early-onset schizophrenia (EOS) patients by applying machine learning algorithms to structural and functional MRI data.

**Methods:** Multimodal MRI data, including T1-weighted MRI (T1w-MRI), diffusion MRI, and resting-state functional MRI (rs-fMRI) data, were collected from 80 antipsychotic-naive first-episode EOS patients and 91 typically developing (TD) controls. The morphometric similarity connectome (MSC), structural connectome (SC), and functional connectome (FC) were separately constructed by using these three modalities. According to these connectivity features, eight brain age estimation models were first trained with the TD group, the best of which was then used to predict brain ages in patients. Individual brain age gaps (BAGs) were assessed as brain ages minus chronological ages.

**Results:** Both the SC and MSC features performed well in brain age estimation, whereas the FC features did not. Compared with the TD controls, the EOS patients presented widened structural BAGs, with opposite trends between childhood and adolescence. These increased absolute BAG scores for EOS patients were positively correlated with the severity of their clinical symptoms.

**Conclusion:** These findings from a multimodal brain age perspective suggest that advanced BAGs exist early in youths with schizophrenia.

## Introduction

The pathologic processes of schizophrenia originate early in brain development (1) and generally manifest as accelerated cortical thinning and synaptic overpruning (2, 3). These processes might lead to structural and functional brain changes that are evident in vivo via magnetic resonance imaging (MRI). Such changes included decreased cortical thickness detected with T1-weighted MRI (T1w-MRI) data (4), reduced fractional anisotropy detected with diffusion tensor imaging (DTI) data (5), and widespread dysconnectivities detected through resting-state functional MRI (rs-fMRI) data (6). Recent MRI-based schizophrenia studies have modelled disease effects from a biological perspective by using data-driven machine learning strategies (7); however, patients’ deviations from typical brain developmental trajectories remain unestablished.

The progress of neurodevelopment can be indexed by the biological age of the brain estimated via information derived from MRI data (8–10). In particular, high-dimensional MRI data for each individual can be transformed into a predicted brain age by machine learning algorithms. The difference between the predicted brain age and the observed chronological age, referred to here as the brain age gap (BAG), was previously considered a prediction error, demonstrating a lack of model accuracy or insufficient data quality. However, the physiology aspect of BAGs has been increasingly reported in recent studies. In the general population, BAG scores vary from one to five years (11–14). BAGs have been shown to be sensitive to cognitive and functional capacities (8). For example, a positive BAG score, which indicates an “older” brain age relative to a person’s chronological age, is related to cognitive deficits (15), early mortality (16), nicotine and alcohol intake (17), and obesity (18). Moreover, advanced brain ages have been associated with several brain diseases, including schizophrenia, bipolar spectrum disorder, and major depression disorder (19). Hence, BAGs are of great neuroscientific and clinical value, and may serve as a reliable indicator of patients’ deviations from healthy brain development trajectories.

Increased BAG scores have been well documented in patients with schizophrenia, with the reported scores varying from +2.6 to +7.8 years across different studies (20–25). Higher BAG scores are related to poorer functioning (19, 26) and clinical outcomes (21, 23) in schizophrenia patients. Moreover, BAG scores gradually increase across the at-risk, recent onset, and recurrent states of schizophrenia (21). A longitudinal study also revealed progressively increased BAG scores during follow-up periods, especially during the first 5 years after the onset of an illness (23). These observed BAG increases are in line with the hypothesis of accelerated brain ageing in patients with schizophrenia (27). In particular, this hypothesis attributes the increased risks of age-related comorbidities (28) and premature mortality in schizophrenia patients (29) to accelerated brain ageing (30), and further suggests that older biological age is the substrate of the progressive decline exhibited by chronic schizophrenia patients. However, increased BAGs have also been found in adolescents and young adults (aged 16-22 years) with schizophrenia (31) and early stage schizophrenia (20). Given that ageing might lie on a continuum with developmental processes that start at birth (32, 33), it is plausible that advanced brain ageing (as quantified by higher BAG scores) could emerge earlier in youths with schizophrenia during brain development.

Schizophrenia patients undergo various changes in their brain structures and functions, but few studies have estimated brain ages via multimodal imaging data. For example, most studies have built brain age estimation models by using single-modal brain structure images, such as T1w-MRI data (7, 16, 19–23) and DTI data (25, 34). Different brain imaging modalities have distinct developmental trajectories and have been suggested to be encoded by diverse sets of genomic loci (35, 36). A recent study revealed different genetic architectures behind the BAG scores derived from T1w-MRI data, DTI data, and rs-fMRI data (37). Specifically, T1w-MRI-based BAG scores are highly genetically heritable and are related to many diseases, conditions, and clinical phenotypes. The genetic architecture of DTI-based BAG scores is strongly correlated with cancer-related traits, autism disorder, and physical measures. Although rs-fMRI-based BAG scores are less genetically heritable than other scores are, their genetic basis still mildly covaries with educational performance and schizophrenia subtypes. Together, all these imaging modalities can serve as reliable and complementary endophenotypes that are close to the underlying aetiology (38). Thus, to better capture the distinct neurobiological facets of developmental deviations, this study estimated patients’ brain ages by using three imaging modalities, including T1w-MRI, DTI, and rs-fMRI data.

In this study, we evaluated early-onset schizophrenia (EOS) patients aged between 7 and 17 years to investigate patients’ deviations from typical brain development trajectories. We summarized high-dimensional neuroimaging data (T1w-MRI, DTI, and rs-fMRI data) as a brain age index by using supervised machine learning algorithms. Specifically, a brain age estimation model was first built by ‘learning’ the relationship between multimodal brain connectomes and chronological ages in a training dataset consisting of a typically developing (TD) population; this model was then used to predict brain ages with patients’ images as its inputs. Next, BAG scores were generated by calculating the differences between the brain-predicted ages and chronological ages. The amplitude of the BAG reflects the extent to which a patient deviates from healthy brain development trajectories. According to previous findings (7), we assumed increased BAGs in EOS patients relative to the TD controls, and this situation was associated with clinical symptoms in patients.

## Materials and methods

### Participants

A total of 199 participants, including 99 antipsychotic-naive first-episode EOS patients and 100 TD controls, were recruited. The EOS patients were selected from the First Hospital of Shanxi Medical University, and the TD controls were collected from the local community through advertisements. All participants were right-handed and aged between 7 and 17 years. The schizophrenia diagnoses followed the criteria outlined in the Structured Clinical Interview for DSM-IV and were confirmed by at least one senior psychiatrist (Y.X.) after a minimum 6-month follow-up period. The psychiatric symptomatology of 71 patients was evaluated via the Positive and Negative Syndrome Scale (PANSS). Subjects were excluded if they had neurological MRI anomalies, electronic/metal implants, or histories of substance abuse. EOS patients were also excluded if they had been suffering from the illness for more than 1 year. TD controls were excluded if they or their first-degree relatives had histories of psychiatric disorders. This study received approval from the Ethics Committee of the First Hospital of Shanxi Medical University. Written consent was obtained from all participants and their parents or legal guardians.

### Data acquisition

MRI data were collected from three imaging modalities via a 3 Tesla MRI scanner (MAGNETOM Verio; Siemens, Germany) at the First Hospital of Shanxi Medical University. T1w-MRI data were acquired using a three-dimensional fast-spoiled gradient echo sequence with the following parameters: repetition time (TR) = 2,300 ms; echo time (TE) = 2.95 ms; flip angle = 90°; matrix = 256 × 240; slice thickness = 1.2 mm (no gap); and voxel size = 0.9375 × 0.9375 × 1.2 mm^3^, with 160 axial slices. DTI data were acquired through a two-dimensional echolJplanar imaging sequence with the following parameters: TR = 6,000 ms; TE = 90 ms; flip angle = 90°; matrix = 128 × 128; number of volumes = 39; slice thickness = 3 mm (3 mm gap); voxel size = 1.875 × 1.875 × 3 mm^3^, with 45 axial slices; b = 1000 s/mm^2^ for 36 diffusion directions; and three b = 0 s/mm^2^ images. Rs-fMRI data were obtained via a two-dimensional echolJplanar imaging sequence with the following parameters: TR = 2,500 ms; TE = 30 ms; flip angle = 90°; matrix = 64 × 64; number of volumes = 212; slice thickness = 3 mm (1 mm gap); and voxel size = 3.75 × 3.75 × 4 mm^3^, with 32 axial slices. During rs-fMRI scanning, all the subjects were instructed to keep their eyes closed, avoid engaging in specific thoughts, maintain a stable head position, and refrain from falling asleep.

### Data preprocessing

#### T1w-MRI data

The T1-weighted anatomical images were preprocessed via the FreeSurfer package (v7.1.0, http://surfer.nmr.mgh.harvard.edu/) (39). The anatomical surface was first obtained from each image in the native space through the recon-all processing techniques, including skull stripping, cortical extraction, the segmentation of cortical white and grey matter, the separation of the hemispheres and subcortical structures (40), and surface reconstruction of the grey/white interface and the pial surface. On the basis of the individual surface and volume templates, five anatomical features were then extracted from each individual’s surface data, including their grey matter volume, surface area, cortical thickness, Gaussian curvature, and mean curvature. During these preprocessing steps, four patients were excluded because of incomplete scans, and two controls were excluded because of poor-quality cortical parcellation effects, resulting in a final T1w-MRI data sample that included 95 EOS patients and 98 TD controls.

#### DTI data

Structural diffusion images were preprocessed via MRtrix3 (https://www.mrtrix.org/) (41), the FMRIB Software Library (FSL, v5.0.9, http://www.fmrib.ox.ac.uk/fsl) (42), and the Advanced Normalization Tool (ANTs, https://stnava.github.io/ANTs/). The individual images were first denoised and corrected for eddy currents and bias fields and were then used to conduct tractography via MRtrix3. By using a unidirectional tracking approach, a total of 5 million streamlines were generated, with a minimum length of 20 mm and a maximum length of 200 mm. The maximum angle between successive steps was set to 45 degrees, and a fractional anisotropy cut-off of 0.06 was applied as previously suggested (43). Each image was then coaligned with its nondiffusion-weighted B0 image and further mapped onto an individualized functional partitioning template transformed from the Montreal Neurological Institute space via ANTs. Nine EOS patients and one control were excluded because of incomplete scans, resulting in a final DTI data sample consisting of 98 TD controls and 90 patients.

#### Rs-fMRI data

The functional images were preprocessed with the CBIG pipeline (https://github.com/ThomasYeoLab/CBIG) based on the FSL (v5.0.9) and FreeSurfer (v7.1.0). The processing steps included removal of the first four volumes, slice-timing, motion correction, boundary-based registration to structural images, covariate regression, and bandpass filtering (0.01–0.08 Hz). Participants with visually severe segmentation faults and intrasubject registration costs exceeding 0.7 were discarded. Volumes with FD > 0.2 mm or voxel-wise differentiated signal variances > 50 were marked as outliers. Censored images were excluded and substituted by the least-squares interpolation of the neighbouring time frames. Participants with mean FD > 0.2 mm or censored volume percentages > 50% were excluded from further analyses. White matter signals, ventricular signals, head motion parameters and temporal derivatives were regressed out to remove spurious noise effects. Thirty patients and nine controls were excluded because of nonideal registrations from the functional images to the anatomical images, resulting in a rs-fMRI data sample consisting of 91 controls and 86 patients.

Taken together, the preprocessed multimodal imaging data included a T1w-MRI sample containing 98 TD controls and 95 EOS patients, a DTI sample consisting of 98 controls and 90 patients, and a rs-fMRI sample including 91 controls and 86 patients. After combining the imaging data for the three modalities, a total of 91 TD controls and 80 demographic-matched EOS patients were further included in the following analyses (see **Table 1** for the demographic data).

**Table 1.**
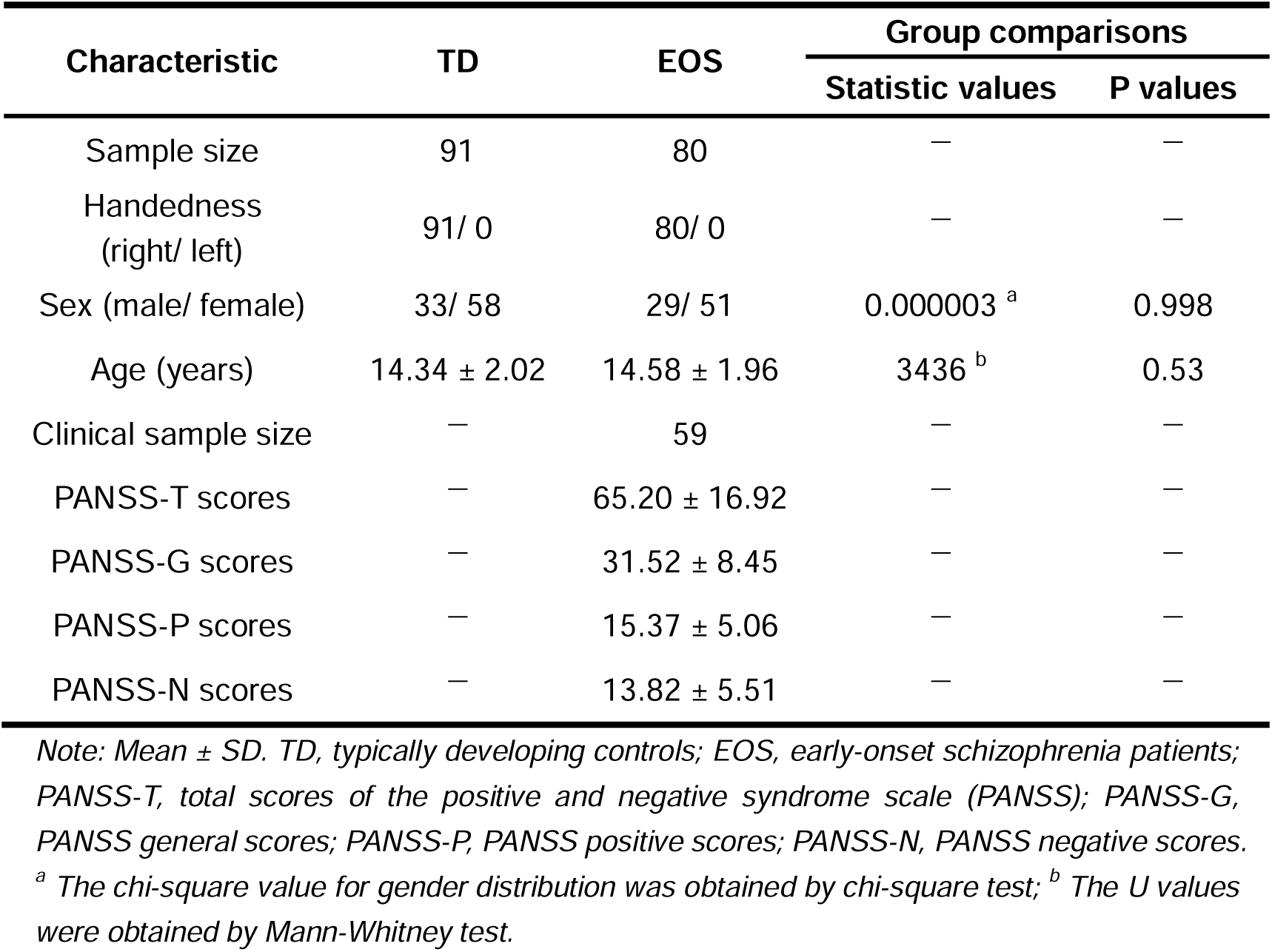
Demographic and clinical characteristics.

### Construction of multimodal connectomes

To construct multimodal connectomes, the cortical surfaces of all the imaging modalities were parcellated into 360 cortical parcels derived from the “Glasser” multimodal parcellation atlas (44), which consists of 180 symmetric cortical parcels per hemisphere. For the T1w-MRI data, parcel-wise morphometric features were first calculated by averaging the vertex-wise feature values within each parcel, creating a five-feature vector for each parcel (grey matter volume, surface area, cortical thickness, Gaussian curvature, and mean curvature). Morphometric similarity scores were then measured by computing Pearson’s correlation coefficient between each pair of morphometric feature vectors. For each subject, we obtained a 360 × 360 morphometric similarity connectome (MSC).

With respect to the preprocessed DTI data, the average fractional anisotropy values of all fibres connecting two regions were first calculated. Next, the reconstructed streamlines were mapped onto 360 cortical parcels. Structural connections were defined by the number of streamlines between two parcels (i.e., the fibre density). After log-transforming the connectivity values to reduce the connectivity strength variance, we obtained a 360 × 360 structural connectome (SC) for each subject.

For the rs-fMRI data, vertex-wise functional images were first downsampled to 360 parcels. Pearson’s correlations were then computed between different pairs of time series consisting of 360 parcels, generating a raw connectivity matrix for each subject. Next, the negative connectivity values of the matrix were set to zero as previously suggested (43). After *z*-transforming connectivity matrix, we obtained a 360 × 360 functional connectome (FC) for each subject.

### Brain age estimations based on multimodal connectomes

We first grouped the 360 cortical parcels into 22 networks comprising geographically contiguous sets of parcels on the basis of their structural and functional characteristics (see **Table S1** for details). A total of 253 connectivity features were extracted for each imaging modality, including 22 within-network connections and 231 between-network connections. These features were then normalized for each modality, resulting in 3 × 253 connectivity features for each subject. To further select age-related features, we performed a cross-validation process entailing recursive feature elimination (45) on these connectivity features. Moreover, to eliminate the impact of head motion, we excluded functional connectivity features that were significantly associated with mean frame directional displacements.

To predict multimodal brain ages, we used eight regression models, including linear support vector regression (SVR), elastic network regression, ridge regression, least absolute shrinkage and selection operator (LASSO) regression, linear regression, decision tree regression, random forest regression (RFR), and k-neighbours regression models. The algorithms were implemented in Python via the scikit-learn package (sklearn, https://scikit-learn.org/) (46). Based on the assumption that brain age was significantly related to chronological age in the normative population (12), the connectivity features selected for the three imaging modalities were defined as input features, and chronological age was used as a predictor. Here, the TD group was used to build the brain age estimation model. Specifically, all 91 controls were randomly split into training (90%) and testing (10%) datasets, and the model parameters were optimized through 5-fold cross-validation: the training set was equally separated into five parts (four for model estimation and one for validation purposes), and this part repeated five times to employ all the parts as the validation set. The model parameters that led to the highest cross-validation accuracy were then applied on the testing set to predict brain ages. The performance of each model was evaluated by comparing the chronological and estimated brain ages via the coefficient of determination (R^2^) and mean absolute error (MAE) metrics. The statistical significance of the correlation between brain age and chronological age was tested via permutation tests (*p_perm_* < 0.05, 5,000 times). The procedure, including model learning and testing, was repeated 50 times, resulting in 50 fitted models. Across eight regression models, the model with the best average performance was selected for the subsequent analyses.

The best model trained on the TD group was used to estimate brain ages in the EOS group. A BAG score was calculated for each subject by using the discrepancy between the predicted brain age and their observed chronological age. Here, a BAG score greater than zero indicated a brain that appeared “older” than the person’s chronological age, whereas a BAG score lower than zero reflected a “younger” brain than expected at a given chronological age. We further assessed the predictive weights of the connectivity features by using feature coefficients in the model.

### Clinical associations with brain age

To investigate the clinical relevance of the brain age deviations exhibited by EOS patients, we conducted correlation analyses between the clinical symptoms and BAG scores. Among the EOS patients, under all three imaging modalities, the clinical symptoms of 59 patients were evaluated via the PANSS. We computed Pearson’s correlations coefficients between the total, positive, and negative PANSS scores and the BAG scores. The outliers in the correlation analyses were identified and removed by means of the bootstrapped Mahalanobis distance measure (47).

## Results

Multimodal imaging data, including T1w-MRI, DTI, and rs-fMRI data, were first downsampled to form a 360-parcel multimodal parcellation atlas (44) and separately used to construct MSC, SC, and FC matrices (**Figure 1A**). These parcel-wise connectivity features were then grouped at the network level (see **Table S1** for the network details), and filtered via a recursive feature elimination approach, generating 36 MSC features, 12 SC features, and 249 FC features (**Figure 1B**). Next, these selected connections were separately used as input features in eight regression models to predict an individual’s brain age (**Figure 1C**). Specifically, the TD group was used to construct a brain age prediction model. On the basis of the assumption that the brain changes as we age and that these changes are consistent across individuals (8), the performance of the models was measured by comparing the predicted brain ages and the chronological ages. Finally, the model with the best performance was used to further estimate the brain ages in the EOS group.

**Figure 1.**
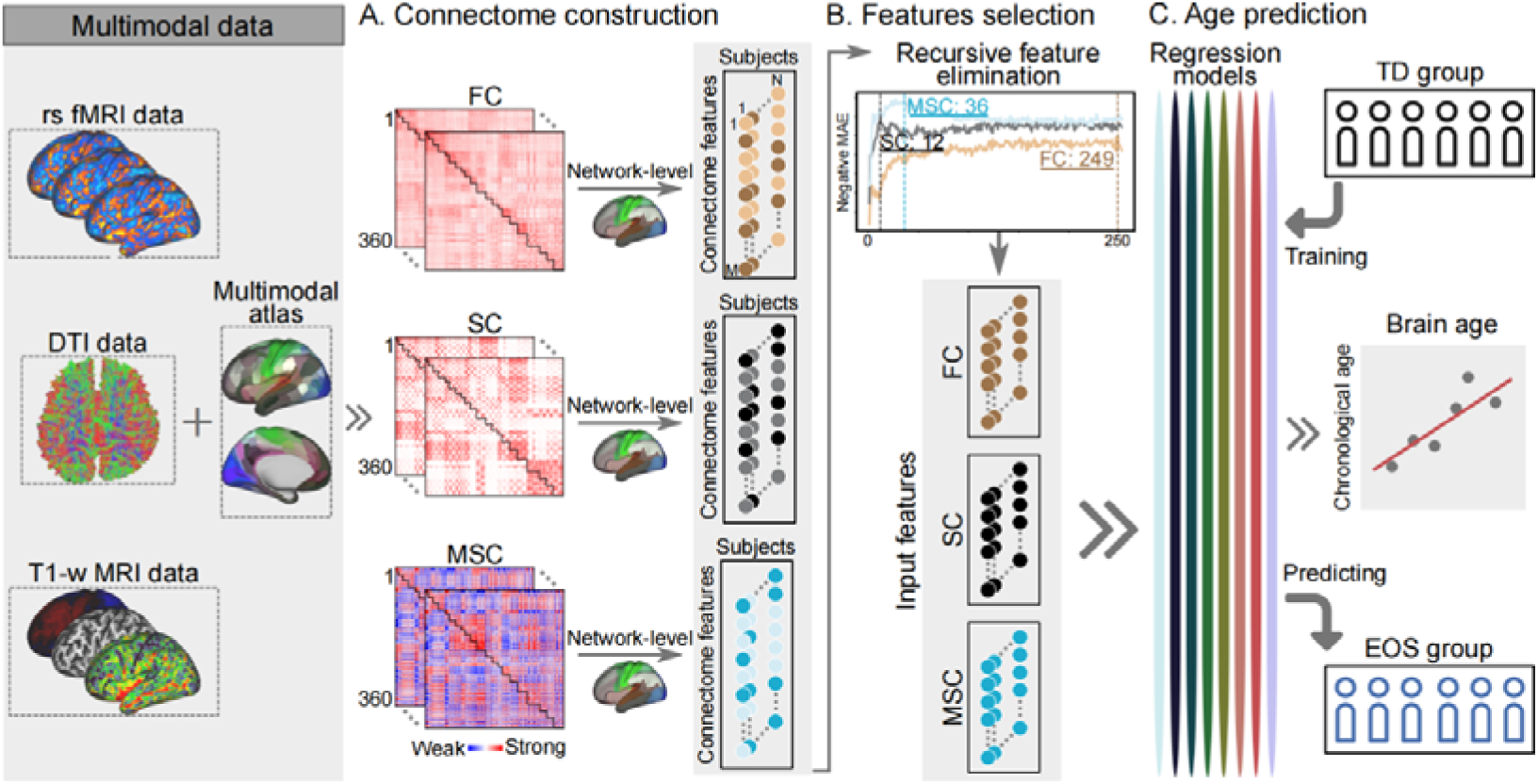
Flowchart of the proposed method. rs-fMRI, resting-state functional MRI; DTI, diffusion tensor imaging; T1w-MRI, T1-weighted MRI; FC, functional connectome; SC, structural connectome; MSC, morphometric similarity connectome; TD typically developing; EOS, early-onset schizophrenia.

### Brain age estimation models

In the TD group, brain ages were separately predicted with eight regression models, including SVR, elastic network regression, ridge regression, LASSO regression, linear regression, decision tree regression, RFR, and k-neighbours regression models (**Figure 2A**). The estimation procedure, including model learning and testing, was repeated 50 times. The performance of the models was evaluated by using the mean values of the R^2^ and the MAE between the brain and chronological ages across 50 fitted models. We found that the RFR model attained the best performance on the basis of the FC, SC, and MSC features (**Table 2**). By testing the statistical significance of the correlation between brain age and chronological age (*p_perm_* < 0.05, 5,000 times), we found that the RFR model also showed the highest correlation relative to the other models (**Table S2**). Therefore, the RFR model was further used to predict brain ages in the EOS group. However, the R^2^ values produced by all the regression models based on the FC features were close to zero, indicating poor model performance. The regression models using all FC, SC, and MSC features achieved good brain age estimation performance for the TD group (**Figure S1**).

**Figure 2.**
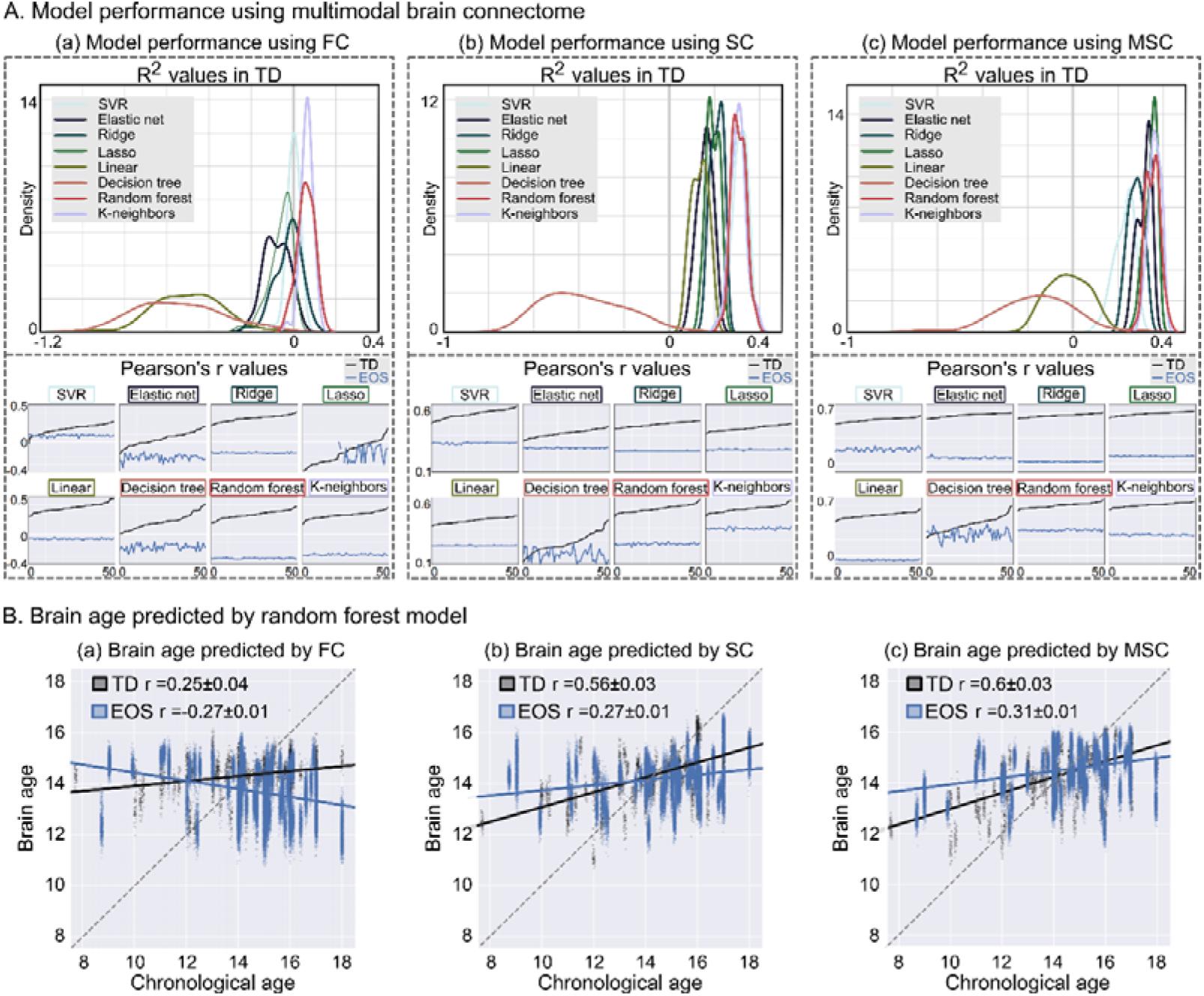
Predicted brain ages for TD and EOS. **(A)** Performance of eight regression models. Eight models, including linear SVR, elastic network, ridge regression, LASSO regression, linear regression, decision tree, random forest, and k-neighbours regression models, are shown in different colours. The prediction procedure was repeated 50 times, and the R^2^ values between the predicted ages and observed chronological ages were used to evaluate the performance of the models. The TD group is represented in black, and the EOS group is represented in blue. **(B)** Correlations between the brain ages and chronological ages. For 50 repeated fitted models, we report the means and standard deviations of the r coefficients. The slope of the grey dashed reference line equals 1.

**Table 2.**
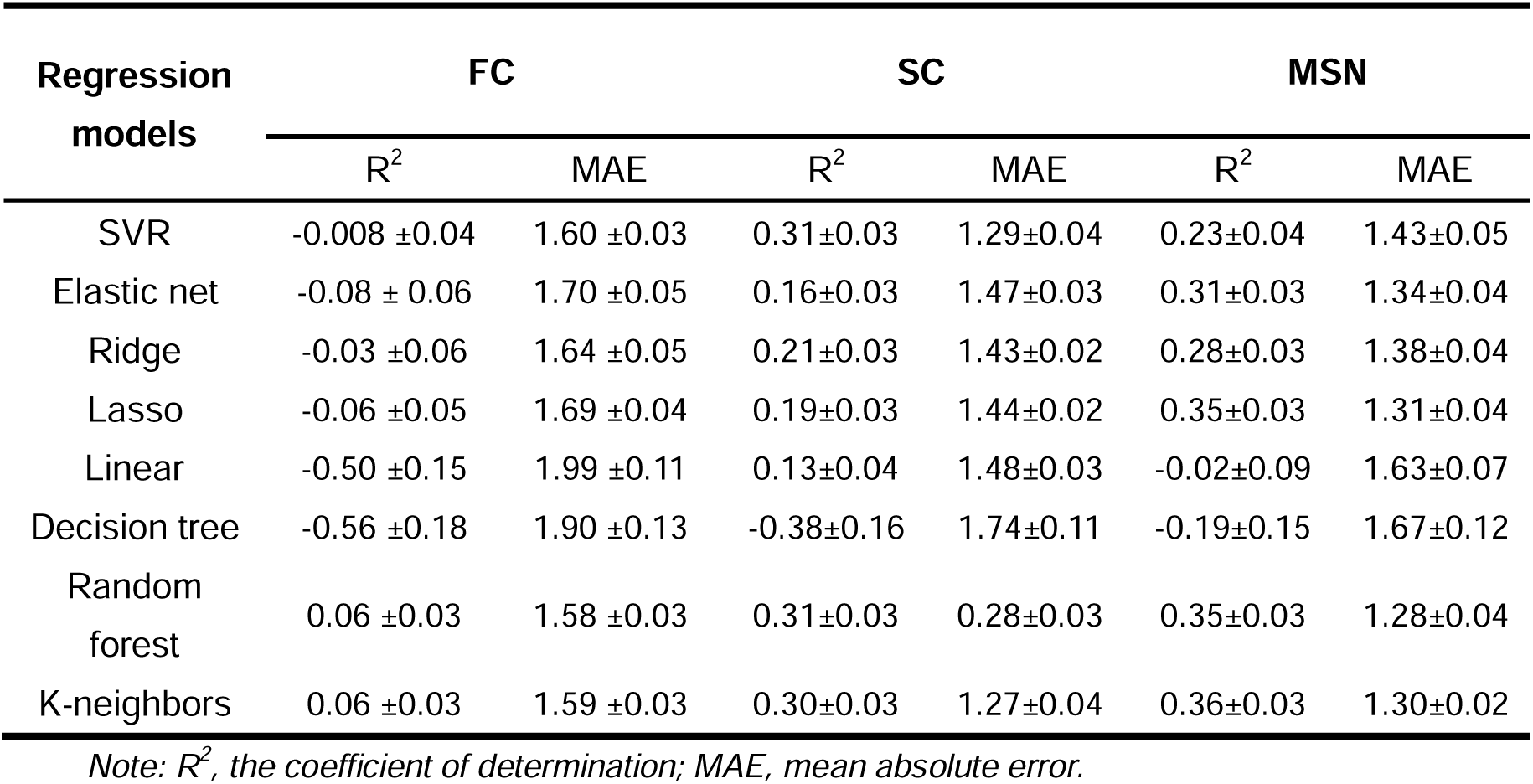
The performance of 8 regression models in TD controls.

Compared with the TD controls, EOS patients presented weaker correlations between their brain ages and chronological ages when FC (*r* = −0.27 ± 0.01; two-sample *t* test, *t_50_* = −82.07, p < 0.0001), SC (*r* = 0.27 ± 0.01; *t_50_* = −66.84, p < 0.0001), or MSC (*r* = 0.31 ± 0.01; *t_50_* = −64.46, *p* < 0.0001) features were used (**Figure 2B**). In particular, individuals with EOS had greater brain ages in children and lower brain ages in adolescents than the TD individuals did, indicating increased BAGs for these patients. From a visual perspective, the fitted lines between the brain and chronological ages were nonlinear for the EOS patients. Thus, we performed an exploratory analysis by fitting the lines via two-order regression models and found that the brain ages of patients with EOS indeed showed a nonlinear relationship with their chronological ages, but this did not hold for the TD group (**Figure S2**).

### BAG scores and clinical relevance

In the SC-based brain age estimation model (**Figure 3A**), predictive features with heavier weights presented hyperconnectivities in EOS patients compared to TD controls, whereas the other features presented hypoconnectivities in EOS. After summing the predictive weights of each network, we found that the premotor network of the sensorimotor cortices presented the highest predictive weight. In the MSC-based brain age estimation model (**Figure 3B**), the major predictive features showed increased morphometric similarity covariances in the patients relative to the controls. The neighbouring visual network exhibited the highest predictive weights. Here, the FC-based prediction model is not shown due to its poor model performance, but its results can still be found in the supplementary materials (**Figure S3**). Briefly, the FC-based results suggested that the dorsolateral prefrontal network showed the highest predictive weights, rather than the unimodal networks that demonstrated the highest weights in the structure-based results.

**Figure 3.**
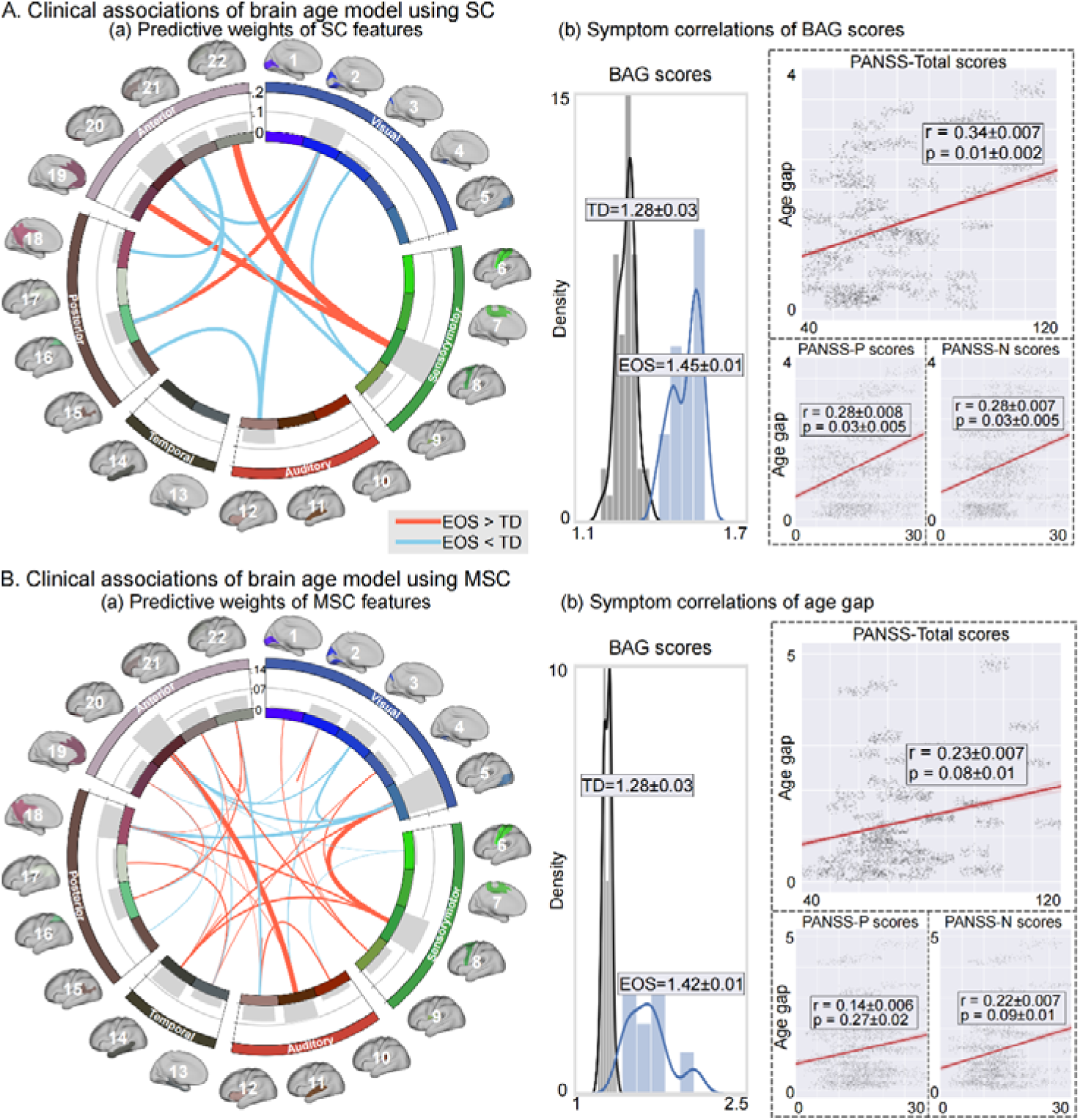
Clinical associations. The order of the 22 networks is the same as that in **Table S1**. The TD group is represented in black, and the EOS group is represented in blue. BAG, brain age gap; PANSS, the Positive and Negative Syndrome Scale; PANSS-P, positive PANSS scores; PANSS-N, negative PANSS scores.

For individuals with EOS, increased BAG scores were found in both prediction models based on the SC features (*t_50_* = 35.51, *p* < 0.0001) and the MSC features (*t_50_* = 26.58, *p* < 0.0001). Moreover, patients’ SC-based BAG scores were positively related to their total PANSS scores (*r* = 0.34 ± 0.007, *p* = 0.01 ± 0.002), positive PANSS scores (*r* = 0.28 ± 0.008, *p* = 0.03 ± 0.008) and negative PANSS scores (*r* = 0.28 ± 0.007, *p* = 0.01 ± 0.002). Here, we removed one outlier from the correlation analysis (see **Figure S4** for correlations without removing outlier points). The MSC-based BAG scores did not significantly correlate with the total PANSS scores (*r* = 0.23 ± 0.07, *p* = 0.08 ± 0.01), positive PANSS scores (*r* = 0.14 ± 0.006, *p* = 0.27 ± 0.02), and negative PANSS scores (*r* = 0.22 ± 0.007, *p* = 0.09 ± 0.01). In the brain age prediction model based on all feature modalities (**Figure S5**), we still found higher BAG scores for the EOS group than for the TD group (*t_50_* = 63.61, p < 0.0001), with a positive association with the negative PANSS scores (*r* = 0.27 ± 0.006, *p* = 0.04 ± 0.005).

## Discussion

This study investigated deviations in the brain ages of EOS patients from the trajectories of TD controls by implementing several machine learning approaches on multimodal imaging data. We found that the RFR model performed best during brain age estimation, and the structural imaging-based prediction models performed better than the functional models did. Patients’ brain ages were greater than their chronological ages during childhood and lower in adolescence. All individuals with EOS showed advanced BAGs relative to the controls, which were reflected by their increased BAG scores. These increased BAG scores were positively related to the severity of the clinical symptoms exhibited by the EOS patients. Overall, this study revealed widened BAGs in youths with EOS via the use of multimodal connectome features, indicating early neurodevelopmental deviations in patients with schizophrenia.

Conceptually, brain age is a measure of biological neurodevelopment that capitalizes on recent neuroscientific advancements to digitize the brain according to multimodal imaging. Although machine learning algorithms can eliminate discrepancies between observed ages and ages predicted on the basis of anatomical measures (48), recent studies have suggested that the differences between predicted brain ages and chronological ages may reflect normal variations. For example, increased brain ages can appear even in relatively healthy older people and are associated with cognitive processing later in life (49). Here, we found that the predicted brain ages were not perfectly aligned with the chronological ages of the individuals in the normative population. Specifically, brain ages are greater than chronological ages in children aged younger than 14 years, but lower than chronological ages in adolescents aged older than 14 years. Mean absolute prediction errors of approximately 1.0–1.7 years from chronological ages have been reported in several studies (12, 31, 50), and these errors were interpreted as delayed or accelerated brain development relative to the actual chronological ages. Moreover, a recent study based on multimodal feature sets consistently suggested increased brain ages in younger children and decreased brain ages in older adolescent participants (51). Except for one explanation concerning the reflection of individual differences in outwardly healthy people, these age-specific differences could also be explained by a bias effect of the utilized regression model (52); i.e., brain ages are overestimated in younger subjects and underestimated in older subjects. In this case, a common practice is to apply a statistical bias correction to the age prediction process. After applying a bias adjustment scheme to reduce the prediction error (53), our estimation models performed better in the normative population than did the strategies with no bias adjustment (**Figure S6**), whereas the stage-specific differences between the predicted brain ages of the patients and controls were retained. Therefore, despite this model error, BAG scores could be used to index the extent to which an individual deviates from age-expected brain development.

In keeping with the findings in chronic schizophrenia patients (7), individuals with EOS also showed increased BAGs relative to those of the healthy controls. However, the widening of the BAGs in adults with schizophrenia was reflected by advanced brain ages relative to their chronological ages (7, 54, 55). Hence, researchers have suggested that accelerated brain ageing in patients with schizophrenia that may be responsible for increased risk of age-related comorbidities and premature mortality (29, 56, 57). Here, we found that the widening of the BAGs was opposite between children and adolescents; i.e., patients’ brain ages were greater than those of the controls during childhood but lower than those of the controls during adolescents. Notably, “younger brain ages” relative to the corresponding chronological ages have also been reported in adolescents with EOS and those at clinical high risk for psychosis (CHR-P) (58). The distinct directions of the widening of the BAG might reflect several age-specific cohort characteristics. Regarding the pathophysiological mechanisms underlying the stage-specific widening of the BAG in EOS patients, both accelerated cortical thinning (4) and excessive synaptic pruning (2) during brain maturation might be involved. Future studies incorporating molecular indices are needed to further investigate the nature of age-specific BAG widening in individuals with EOS during their development.

Compared with that in chronic schizophrenia patients, the gap in EOS was smaller. The BAG distribution varied across mental disorders such as bipolar disorder and major depression, among which schizophrenia had the widest gap (19). Moreover, the effect of increased BAGs gradually decreased across individuals with EOS, individuals with CHR-P converted to a psychotic disorder, and CHR-P nonconverters (58). Thus, it is reasonable to infer that the lesser degree of impact could be explained by the different stages of disease progression. Nevertheless, we found that advanced BAG scores estimated according to the structural features of grey matter were associated with the severity of clinical symptoms. Taking all the models based on different features together, these clinical associations were more obvious with negative symptoms in the EOS patients. Consistently, it has been widely reported that a greater BAG is positively related to more severe negative symptoms (19, 21, 54). Therefore, these findings suggest that individual brain age differences between individuals with schizophrenia and those with typical developmental trajectories might serve as objective neuroimage marker for clinical assessments. However, in addition to diagnostic effects, many other factors could also contribute to these BAGs, such as stress (59) and obesity (60). Future studies are recommended to examine the potential influencing factors that contribute to patients’ BAGs.

The brain age estimation model constructed on the basis of SC and MSC features performed better than the FC-based model did. Indeed, most previous studies revealed advanced BAGs in schizophrenia patients through the use of the structural features of the white matter or grey matter in the brain. For example, two large-scale collaborative studies have reported increased brain ages in patients with schizophrenia derived from structural T1-weighted MRI (7, 19), and another recent schizophrenia study conducted across 10 data cohorts also reported a moderate increase in brain age via DTI data (34). In addition, increased anatomical BAGs were also reported to be related to psychopathology in children and adolescents (61, 62). Although fMRI data have been used for brain age prediction in patients with schizophrenia, functional models are less robust than structural models in terms of capturing brain maturation patterns (31). Compared with functional brain age, anatomical brain age during neurodevelopment tends to be more stable and heritable (63). In our brain age prediction model including the structural features of both grey matter and white matter, the visual and sensorimotor cortices separately made the greatest contributions. This finding was consistent with a recent study on estimating the brain ages of children and adolescents, which suggested that the most important connections for age prediction were related to the sensorimotor and visual areas (64). Combining these results, these unimodal networks might be central aspects of brain maturation. Further longitudinal studies are needed to investigate the age-related changes exhibited by these predictive features, which might aid in the development of neural interventions for patients with neurodevelopmental disorders, including schizophrenia.

## Conclusions

This study revealed an increased multimodal brain gap in EOS patients relative to TD controls, indicating early neurodevelopment deviations from the typical developmental trajectory in schizophrenia patients. Moreover, greater neurodevelopmental deviations are related to more severe clinical symptoms in patients with EOS. Broadly, these findings, derived from a neurodevelopmental perspective, provide new insight into the accelerate ageing hypothesis for patients with schizophrenia.

## Data and code availability

Multimodal connectomes and brain age in TD and EOS and other data supporting the findings of this study are available at https://github.com/Yun-Shuang/Brain-age-in-EOS. Custom code was also made publicly available under https://github.com/Yun-Shuang/Brain-age-in-EOS. Visualizations were based on circos https://circos.ca/ (65), seaborn https://seaborn.pydata.org/ (66) and workbench (https://www.humanconnectome.org/software/connectome-workbench) combined with ColorBrewer (https://github.com/scottclowe/cbrewer2).

## Acknowledgements

We are grateful to all the participants and their guardians in this study. We thank American Journal Experts (https://www.aje.com) for editing this manuscript. This work was supported by STI 2030—Major Projects 2022ZD0208900, the National Natural Science Foundation of China (62403105, 62333003, 62036003, 82121003, 62373079, 62073058), Medical-Engineering Cooperation Funds from University of Electronic Science and Technology of China (ZYGX2021YGLH201). Y-S.F. was also funded by the China Postdoctoral Science Foundation (2023M740524) and Sichuan Province Innovative Talent Funding Project for Postdoctoral Fellows.

