## Supplement for "Neurodevelopmental deviations in schizophrenia: Evidences from multimodal connectome-based brain ages"

**Tables**

**Table S1.** A multimodal parcellation of cerebral cortex from “Glasser” atlas.

| **Order** | **Network name** | **Left brain coordinates** | | | **Right brain coordinates** | | |
| --- | --- | --- | --- | --- | --- | --- | --- |
|  |  | *x* | *y* | z | x | y | z |
| *Early and intermediate visual cortex* | | | | | | | |
| 1 | Primary visual | 100.49 | 100.49 | 41.14 | 78.06 | 44.54 | 74.33 |
| 2 | Early visual | 108.24 | 108.24 | 42.27 | 71.33 | 44.29 | 76.13 |
| 3 | Dorsal stream visual | 110.07 | 110.07 | 41.96 | 68.71 | 45.22 | 105.38 |
| 4 | Ventral stream visual | 126.84 | 126.84 | 66.78 | 56.71 | 69.79 | 56.17 |
| 5 | MT+ complex and neighboring visual | 136.23 | 136.23 | 52.23 | 42.65 | 56.44 | 73.04 |
| The sensorimotor cortex | | | | | | | |
| 6 | Somatosensory and motor | 127.33 | 127.33 | 101.79 | 53.41 | 104.72 | 124.66 |
| 7 | Paracentral lobular and mid-cingulate | 101.76 | 101.76 | 112.64 | 78.91 | 109.80 | 131.37 |
| 8 | Premotor | 132.33 | 132.33 | 123.28 | 47.08 | 126.89 | 113.35 |
| 9 | Posterior opercular | 143.20 | 143.20 | 114.13 | 38.01 | 119.49 | 83.85 |
| Auditory cortex | | | | | | | |
| 10 | Early auditory | 138.22 | 138.22 | 99.71 | 41.99 | 103.12 | 83.70 |
| 11 | Auditory association | 147.24 | 147.24 | 113.68 | 33.80 | 116.04 | 64.00 |
| 12 | Insular and frontal opercular | 128.90 | 128.90 | 131.26 | 52.83 | 132.81 | 69.33 |
| The rest of the temporal cortex | | | | | | | |
| 13 | Medial temporal | 120.31 | 120.31 | 105.45 | 61.14 | 106.56 | 45.96 |
| 14 | Lateral temporal | 142.79 | 142.79 | 106.25 | 40.16 | 110.62 | 44.61 |
| The rest of the posterior cortex | | | | | | | |
| 15 | Temporo-Parieto-Occipital junction | 147.04 | 147.04 | 72.88 | 33.14 | 79.59 | 89.01 |
| 16 | Superior parietal | 114.57 | 114.57 | 68.68 | 65.16 | 69.39 | 128.20 |
| 17 | Inferior parietal | 139.11 | 139.11 | 68.90 | 40.03 | 74.16 | 108.93 |
| 18 | Posterior cingulate | 98.67 | 98.67 | 72.09 | 81.52 | 73.62 | 103.01 |
| The rest of anterior cortex | | | | | | | |
| 19 | Anterior cingulate and medial prefrontal | 97.13 | 97.13 | 162.09 | 84.50 | 160.69 | 86.92 |
| 20 | Orbital and polar frontal | 111.24 | 111.24 | 172.17 | 71.21 | 173.27 | 57.83 |
| 21 | Inferior frontal | 136.79 | 136.79 | 155.95 | 44.28 | 159.79 | 73.77 |
| 22 | Dorsolateral prefrontal | 120.03 | 120.03 | 158.70 | 61.19 | 161.31 | 109.05 |

**Table S2.** Significant times of r values between brain and chronological ages in the TD and EOS groups.

| **Regression models** | **FC**  **(Times)** | | **SC**  **(Times)** | | **MSN**  **(Times)** | |
| --- | --- | --- | --- | --- | --- | --- |
|  | TD | EOS | TD | EOS | TD | EOS |
| SVR | 14 | 0 | 50 | 50 | 50 | 0 |
| Elastic net | 0 | 19 | 50 | 50 | 50 | 0 |
| Ridge | 49 | 0 | 50 | 50 | 50 | 0 |
| Lasso | 21 | 32 | 50 | 50 | 50 | 0 |
| Linear | 50 | 0 | 50 | 50 | 50 | 0 |
| Decision tree | 13 | 8 | 41 | 11 | 50 | 34 |
| Random forest | 44 | 50 | 50 | 50 | 50 | 50 |
| K-neighbors | 44 | 50 | 50 | 50 | 50 | 50 |

*Note: Total repeat times are 50 (p < 0.05, permutation test with 5000 times). SVR, support vector regression.*

**Figures**

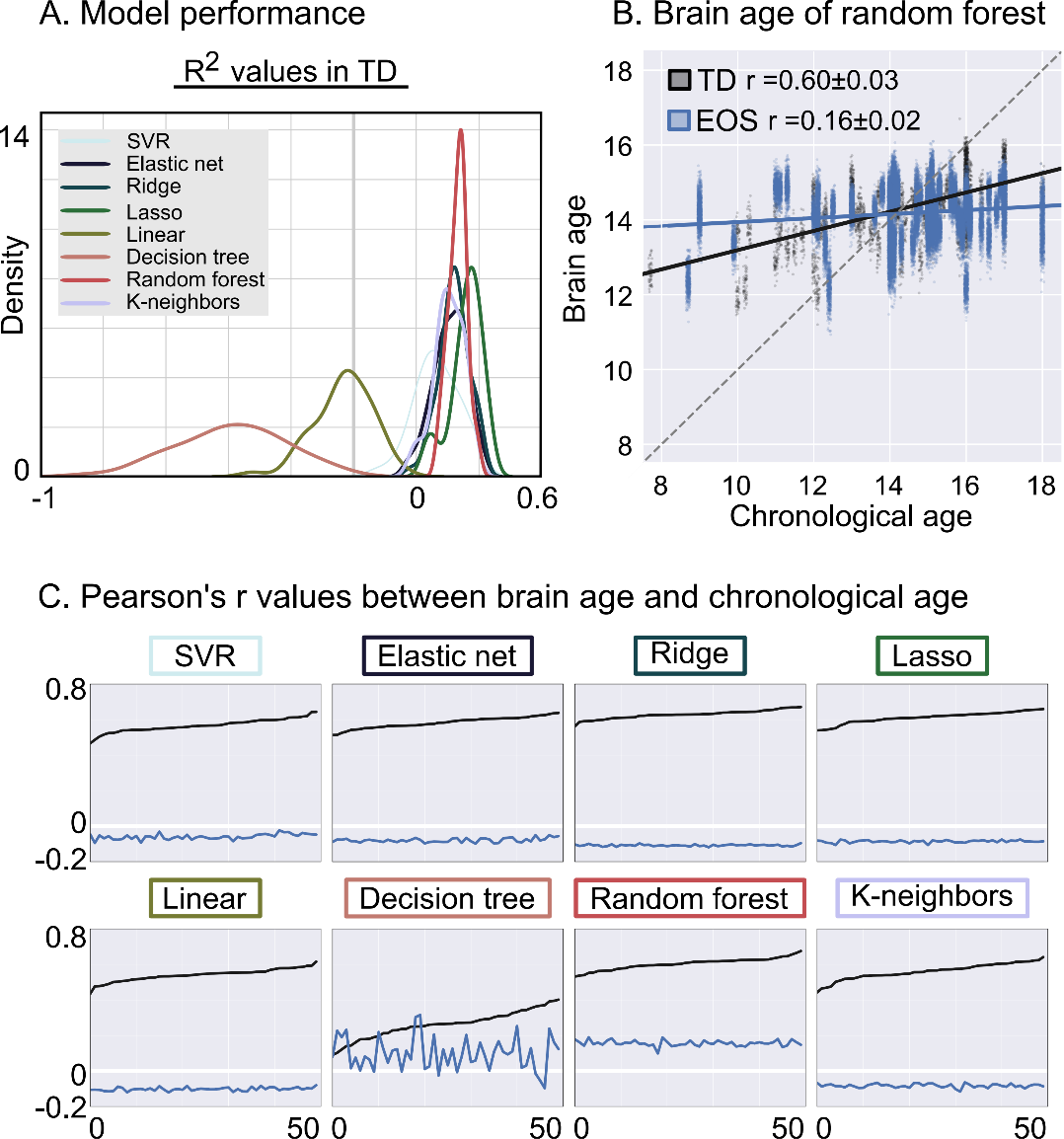

**Figure S1. Predicting brain ages by using all connectomes.** Compared with the TD group, EOS patients showed lower correlation between brain and chronological age by using the FC features (two-sample *t* test, *t_50_* = -87.62, *p* < 0.0001).

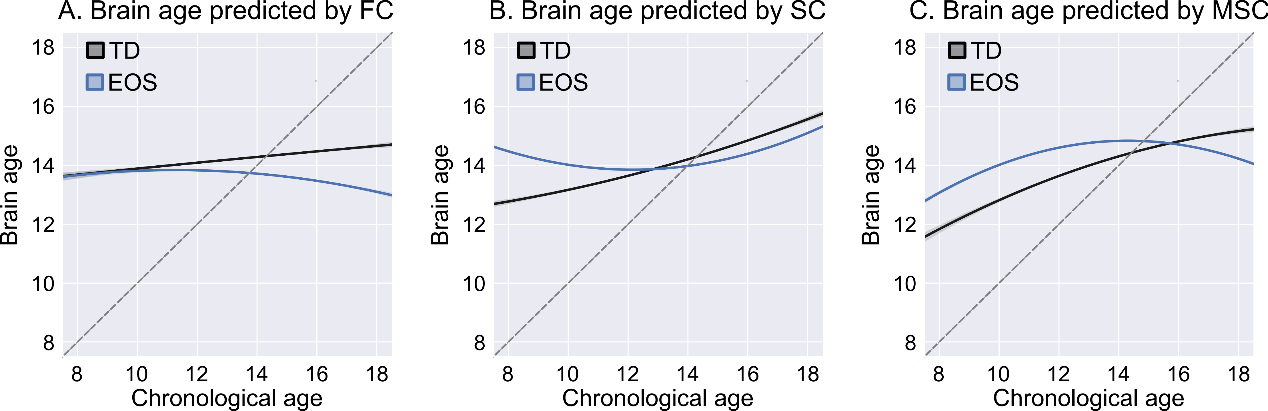

**Figure S2. Two-order regression fit model for brain ages and chronological ages.**

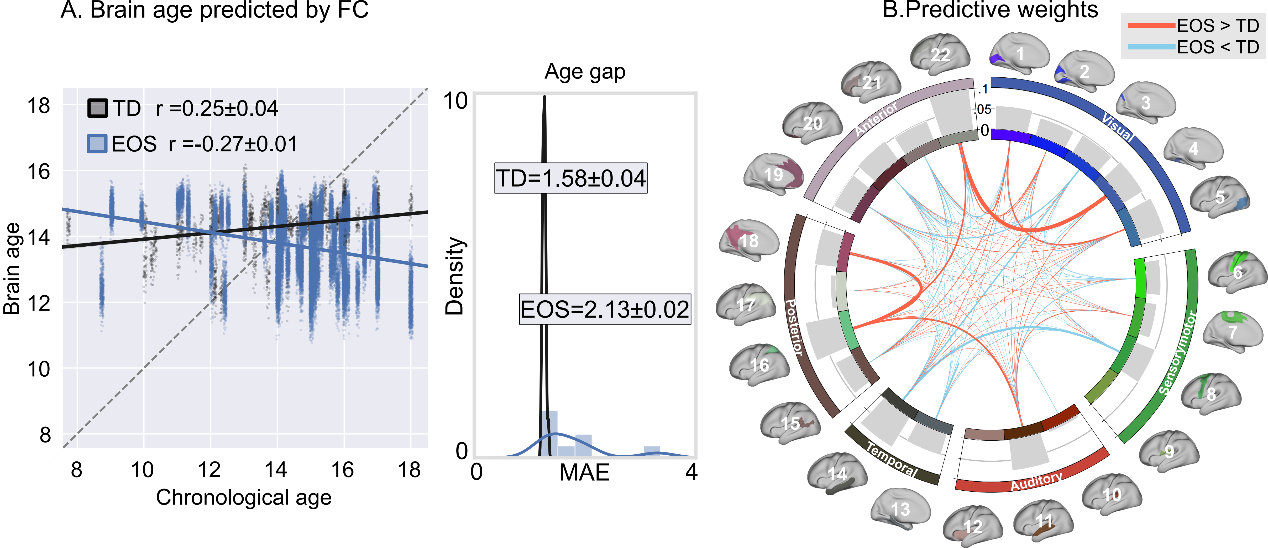

**Figure S3. Brain ages predicted by using the FC features.** Patients had larger age gap (*t_50_* = 110.55, *p* < 0.0001) relative to controls.

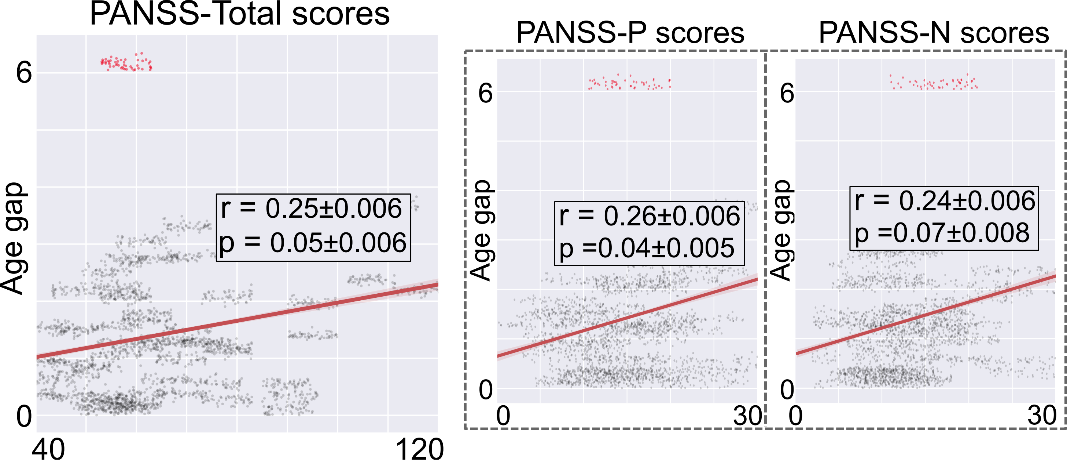

**Figure S4. Clinical correlations of bran age model using SC without deleting outlier.** Absolute age gap in EOS patients showed positive correlation trends with total PANSS scores, positive PANSS scores, and negative PANSS scores. Outliers are shown in red.

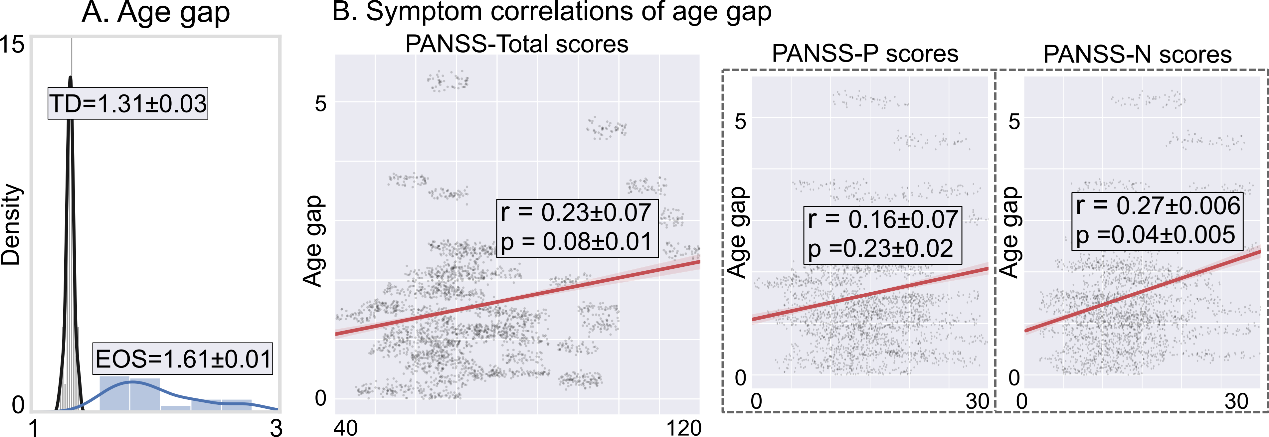

**Figure S5. Clinical correlations of bran age model using all connectomes.**

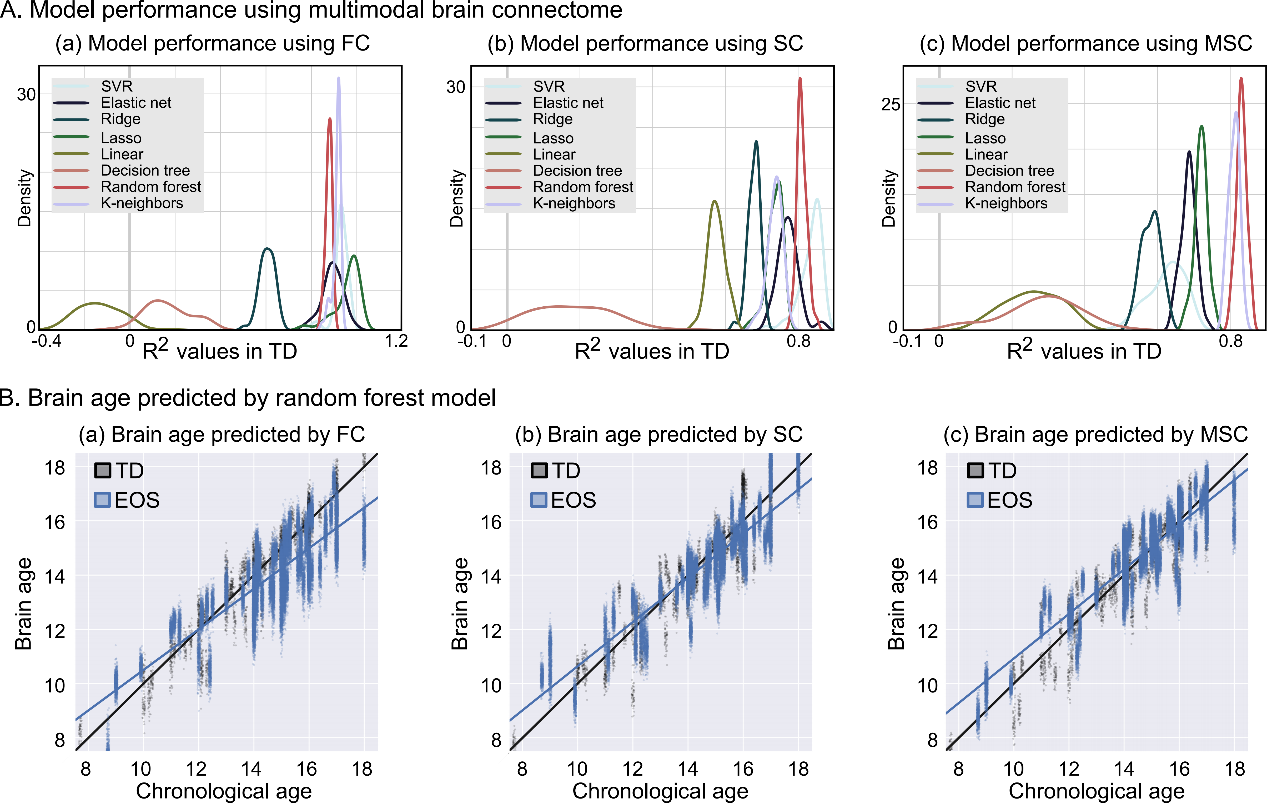

**Figure S6. Brain age estimation models by removing age-related bias effect.**
